# AlphaPeptTools: scverse-native analysis of mass spectrometry-based proteomics

**DOI:** 10.64898/2026.09.11.750886

**Authors:** Vincenth Brennsteiner, Lucas Diedrich, Shani Ben-Moshe, Magnus Schwörer, Georg Wallmann, Wen-Feng Zeng, scverse proteomics consortium, Fabian J Theis, Matthias Mann

**Affiliations:** Department of Proteomics and Signal Transduction, Max Planck Institute of Biochemistry, Martinsried, Germany; Institute of Computational Biology, Helmholtz Center Munich, Munich, Germany; Mikaela Koutrouli, Jose Nimo, Malte Kuehl, Sergio Marco Salas, Matthias Meyer-Bender, Ilan Gold, Marvin Thielert, Laurent Gatto

## Abstract

Mass spectrometry (MS)-based proteomics now routinely profiles proteomes at scale, but extracting biological insight and integrating complementary modalities remains challenging. Here, we present AlphaPeptTools, a Python package for the analysis of MS-proteomics data built on the AnnData data structure of the scverse. AlphaPeptTools implements scalable and modular strategies for MS-data ingestion, quality control, preprocessing, and statistical analysis while seamlessly integrating with the scverse ecosystem, unlocking multi-level, spatial, and multimodal analyses.

Code Availability: https://github.com/MannLabs/AlphaPeptTools.git

## Main

Mass spectrometry (MS)-based proteomics has become a foundational tool of basic and clinical research^1^. Continuous technological innovations yield ever greater proteomic depth in low-input and even single-cell experiments^2–4^. Large-scale studies^5,6^, integration with additional modalities^7,8^, and spatially resolved experiments^9^ are now feasible. Computational tools for MS-proteomics must keep pace on four fronts: efficient storage, standardized analysis, cross-omics integration and interoperability with the open-source ecosystem.

Perseus^10^ and subsequent packages made proteomics analysis accessible to non-programmers; however, their closed nature and graphical user interfaces constrain custom analyses and the addition of new data modalities. R packages such as MSstats^11,12^ or scp^13^ offer robust methods for the statistical analysis of MS-proteomics data but are outside the Python ecosystem where much of single-cell and multi-omics analysis now happens. Clearly, there is a need for standardization, which would allow composing modular workflows and integrating new modalities, without dealing with diverging conventions and source data formats.

Notably, many of these limitations have already been solved in single-cell transcriptomics: the AnnData data structure^14,15^ has been highly successful in standardizing computational analysis, laying the foundation for a rich ecosystem of interoperable and extendable tools called the *scverse*^16^. Recently, several proteomics efforts have utilized AnnData^17–19^ for specific applications, demonstrating the value of scverse data structures and ergonomics in this setting. What is missing is a general-purpose package for MS-proteomics: one that accommodates the unique properties of these data and opens them to integration with complementary omics modalities.

We therefore developed AlphaPeptTools, an open-source Python package that brings MS-proteomics data into AnnData regardless of the search engine that produced it, and with it into the scverse ecosystem (Figure 1A). It reads the quantification output of all major data-independent and data-dependent acquisition (DIA/DDA) search engines (currently 11), in both tabular and binary formats, into AnnData (Supplementary Table 1). We automatically map search engine-specific conventions like column naming, feature representation, and the encoding of post-translational modifications to a unified vocabulary^20^ (Methods). Users can load a single proteomics feature level (precursor, peptide, protein group, or gene) into AnnData, and also connect multiple feature levels with MuData^21^, a collection of AnnData objects. For the latter case, we here introduce the MuLink package, an extension to MuData that encodes the hierarchical structure of proteomics data into a directed acyclic graph, making it possible to query and filter across feature levels (Methods). We expect the general principles implemented in MuLink to improve modeling of related features in many contexts outside of proteomics. Storing search engine peptide-spectrum-match outputs in AnnData markedly reduces on-disk file size across feature levels through reduced redundancy and removal of uninformative search-engine metadata (Figure 1E), supporting the efficient distribution, storage, and analysis of MS-proteomics data.

**Figure 1:**
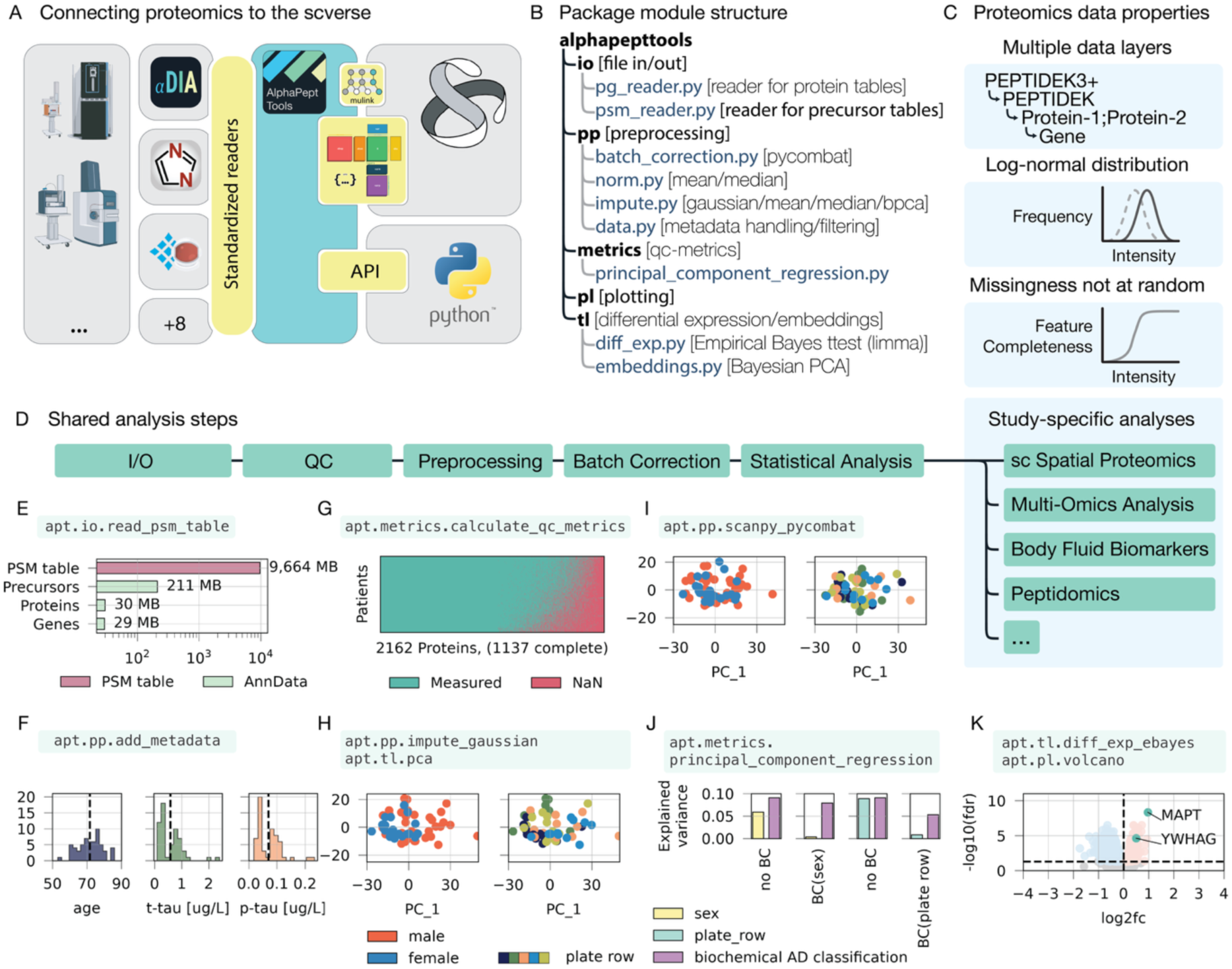
AlphaPeptTools integrates proteomics into the wider-omics and scientific Python ecosystem. **A** The wider Omics-ecosystem with AlphaPeptTools at the intersection between proteomics and the scverse, as well as the general scientific Python stack. **B** AlphaPeptTools package modules with representative functions highlighted. **C** Special features of proteomics data compared to other-omics data: Composition of hierarchical data layers, assumed log-normal distribution, missingness not-at-random (MNAR) with no natural zero. **D** Divergence of different proteomics applications from a shared preprocessing stem. **E** Disk space requirements of a full DIA-NN search engine peptide spectrum match table compared to different data-level AnnData pivots (log-scale). **F** Selection of metadata variable distributions from the Alzheimer’s Disease (AD) dataset, visualized with AlphaPeptTools. **G** Missing vs. measured values in the AD dataset, visualized as a heatmap of missing (red) and measured (green) datapoints, where rows are samples and columns are features. **H** PCA of the AD dataset colored by sex (left) and plate-row (right). **I** Same PCA as H of the AD dataset after batch-correction for sex (left) and plate-row (right). **J** Explained PCA variance of the AD dataset pre and post batch correction for sex or plate-row, with sex, plate-row and AD-classification individually resolved. **K** Differential expression results between AD and control samples, with MAPT and YWHAG genes highlighted.

Building on this unified representation, AlphaPeptTools implements a full MS-proteomics data analysis pipeline, spanning quality control, normalization, imputation, batch correction, statistical testing, and visualization (Figure 1B), adapted to the unique properties of proteomics data (Figure 1C, Methods). Each step consists of unit-tested, modular functions so that alternative analysis strategies can be quickly explored and applied to diverse proteomics applications, as illustrated below (Figure 1D). Preprocessing functions are accompanied by diagnostic plots and quantitative metrics that allow users to choose a suitable analysis strategy. AlphaPeptTools is explicitly designed for LLM-supported analysis: its documentation and worked example studies are written to provide guidance for users as well as serve as LLM-context, so that a coding assistant can compose a complete workflow from standardized, unit-tested building blocks, which the user then inspects and extends with custom code where needed.

We demonstrate AlphaPeptTools on four datasets spanning standard analysis, multi-level feature hierarchies, spatial proteomics and multi-omics integration. The first illustrates how the package supports analysis decisions: a cerebrospinal fluid (CSF) DIA-proteomics dataset from an Alzheimer’s disease (AD) biomarker study^22^. In this example, the PCA function of the *tools* module (*tl.pca*) identifies the biological covariate sex as separating in the first component (Figure 1H, left panel). We apply ComBat batch correction^15,23^ for sex (*pp.scanpy_combat*, Figure 1I), and quantify the variance attributable to that covariate via a one-line call to principal component regression^24,25^ (Figure 1J). Correcting for sex reduces its explained variance from 5.9% to 0.4% while retaining most of the AD signal (9.1% to 7.9%). In contrast, correcting for the technical variable “plate_row” (Figure 1H,I, right panels) removed nearly half of the AD-associated variance (9.1% to 5.3%, Figure 1J), highlighting how batch corrections can incur a substantial loss of biological signal. By making this tradeoff transparent and quickly explorable, users can make informed batch-correction decisions on the spot, rather than through trial and error in the subsequent pipeline. Finally, empirical Bayes moderated differential expression (*tl.diff_exp_ebayes*) analysis between AD and baseline, controlling for the identified covariate sex, recovers the known marker Tau (MAPT) as the most differentially expressed protein (Figure 1K).

The analysis pipeline extends naturally from protein-centric analyses to multi-level proteomics features. In a PEptide Local Stability Assay (PELSA) dataset^26^, treatment with the pan-kinase inhibitor staurosporine locally shields a protein from proteolysis, so that peptides proximal to the binding site in the target protein drop in abundance. From a DIA-NN search engine report^27^, AlphaPeptTools links all quantitative feature levels in a queryable MuLink object (*io.mulink_from_anndatas;* Figure 2A) and runs differential expression^28,29^ analyses across all feature levels (*tl.diff_exp_ebayes*). For the kinase MAP3K7, neither the gene nor its two isoforms are differentially regulated, but querying for its ten quantified peptide precursors (*query.ancestors*) shows that one proximal to the ligand binding site is strongly downregulated in line with the local shielding effect (Figure 2A). MuLink’s hierarchy therefore resolves regulation that protein-level analysis alone would miss and does so in a single query rather than a bespoke filtering operation across separately parsed feature levels.

**Figure 2:**
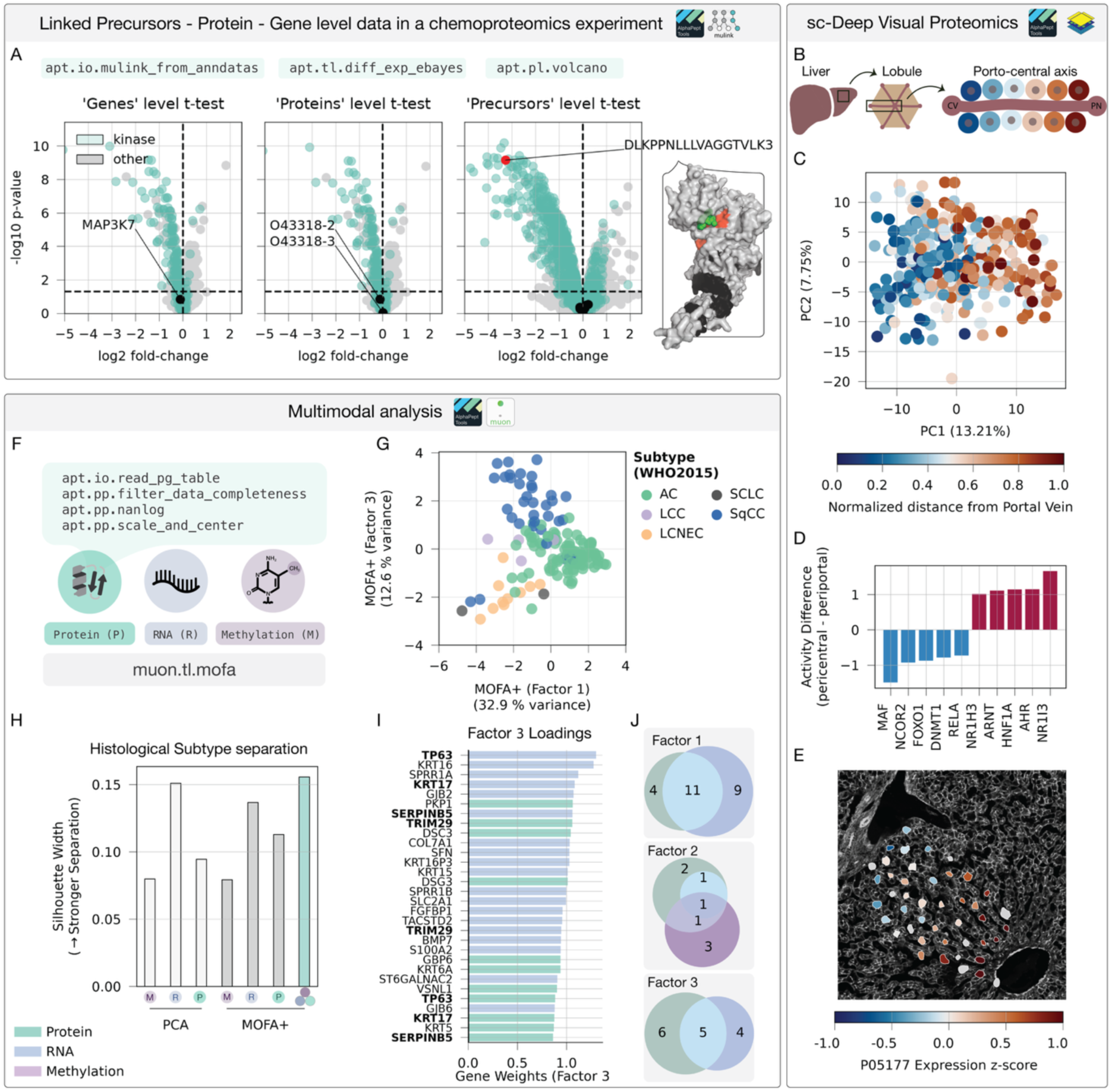
AlphaPeptTools streamlines downstream analysis across MS-proteomics-applications. **A** Differential expression results of a PELSA experiment of pan-kinase inhibitor staurosporine treated cell lysate vs. DMSO-control with kinase-related features (green). A Boltz-2 predicted structure of MAP3K7 (right) with highlighted precursors (red = regulated, black = non-regulated) and staurosporine (green). **B** Hepatocyte differentiation axis in single-cell DVP experiment. **C** PCA colored by localization across periportal-central vein axis. **D** Most-differentially active transcription factors along liver zonation axis in healthy donors, represented as absolute difference between spatial bin 1 and spatial bin 5. **E** Overlay of proteomics features with spatial image of one donor via SpatialData **F** Multimodal integration of DDA MS-proteomics data with RNA- and methylation microarray data **G** MOFA+ embeddings of dimension 1 and dimension 3 of the full clinical NSCLC cohort using three modalities, colored by histological subtype. **H** Separation of histological subtypes with >5 samples in factor models as measured by the silhouette width for single-modalities (PCA/MOFA+ factor models) or joint MOFA+ model of all modalities. **I** Factor 3 loadings showing 30 top-weighted features with protein-features (blue), RNA-features (green), and methylation features (purple, no features in the top-weighted loadings). **J** Overlap of significantly enriched (GSEA, FDR < 0.01) Hallmark genesets for protein (green), RNA (blue), and methylation (purple) weights of the MOFA+ factors 1, 2, 3.

AlphaPeptTools also allows users to leverage existing infrastructure to extend standard proteomics analyses, for example to integrate spatial information. We illustrate this on a spatially resolved single-cell proteomics dataset^30^ in which Deep Visual Proteomics (DVP)^31^ profiled individual human hepatocytes along the porto-central liver axis (Figure 2B). AlphaPeptTools quickly reproduces the published workflow and recapitulates the expected zonation pattern in the first principal component (Figure 2C, Methods). Going beyond our original analysis, we apply decoupler^32^, an scverse package for activity inference, directly to the proteomics AnnData object to estimate transcription factor activities along the spatial axis. This analysis identifies NR1I3 as the most differentially active transcription factor between periportal and central vein (Figure 2D), consistent with its recently reported role in inducing central liver programs^33^. Separately, we bind the proteomics AnnData object directly to the available immunofluorescence images via the scverse-core package SpatialData^34^. This allows the straightforward visualization of spatial protein zonation by overlaying protein abundances on the original tissue, in contrast to the original, extensive custom code (Methods, Figure 2E). In both cases, the shared data structure enables seamless integration of proteomics data with existing functionality from the scverse ecosystem.

Finally, to demonstrate how AlphaPeptTools makes proteomics data accessible for multimodal analyses, we integrate proteomics measurements of non-small cell lung cancer (NSCLC) patients^35^ with partially matched RNA-microarray^36^ and methylation-microarray^37^ data. After reproducing the original DDA-TMT preprocessing workflow with AlphaPeptTools (Figure 2F, Methods), we join AnnData objects for all three modalities into one MuData instance. Whereas the original studies related the modalities post hoc, the shared container allows us to model them jointly with the multi-omics factor model MOFA+^38^ through muon^21^, the scverse framework for multimodal data. Factor 3 of the MOFA+ model separates the three major histological subtypes (Figure 2G): squamous cell carcinoma (SqCC), adenocarcinoma (AC), and large cell neuroendocrine carcinoma (LCNEC), matching the subtype separation of the individual modalities alone (silhouette width, Figure 2H). Among Factor 3’s loadings, TP63, an established immunohistochemical marker of SqCC^39^, carries the highest RNA loading and is among the top 10 weighted protein features (Figure 2I). Gene set enrichment analysis^32,40^ of the top three factor loadings recovers both shared and modality-specific pathways (Figure 2J), showing that the modalities contribute complementary rather than redundant information.

In summary, AlphaPeptTools harmonizes proteomics data into a unified framework for preprocessing, visualization and analysis, tailored to its unique data properties. By directly integrating additional modalities via the scverse ecosystem and favoring modular and composable primitives over end-to-end pipelines, the package supports a wide range of proteomics, spatial, and multimodal analyses. For scverse users, proteomics becomes another accessible modality, with no custom infrastructure to build. For proteomics users, common data analysis tasks no longer need reimplementing, and integration of specialized software tools becomes straightforward. More broadly, by building and extending on established data structures in the scverse community, we hope to ensure AlphaPeptTools’ long-term maintainability while catalyzing the development of novel applications that bridge proteomics with adjacent omics modalities and computational ecosystems.

## Methods

### Data structure and reader

AlphaPeptTools builds directly on AnnData, the standard data structure of the scverse ecosystem. AnnData aligns quantitative measurements, stored in the .X and .layers attributes, with observation- and feature-level metadata in the .obs and .var attributes, respectively. Quantitative MS-proteomics data is commonly reported in two formats: Long-format peptide spectrum match (PSM) tables, containing the full search engine output including precursor, protein and gene-level intensities, and wide-format protein group tables, containing aggregated protein-group quantities. AlphaPeptTools ingests both formats through the reader functions *io.read_psm_table* and *io.read_pg_table*, supporting all common DIA and DDA search engines and file formats (tsv, csv, parquet, hdf5) (Supplementary Table 1). We implemented ingestion and standardization of these reports in our infrastructure package alphabase^20^, which sets column names to a common vocabulary and unifies features (e.g. retention time), as well as the representation of post-translational modifications. AlphaPeptTools’ IO module converts this standardized representation into a wide-format AnnData object, with sample and feature information in the .obs and .var attributes, respectively. Users can select feature levels (proteins, genes, peptides, precursors), quantification methods (e.g. raw, LFQ, iBAQ), and filtering criteria such as FDR thresholds, as well as metadata to be retained, through a consistent interface. Once loaded, AnnData objects can be linked into a single MuData object (*io.mulink_from_anndatas*). Our package MuLink formalizes relationships between features as a directed acyclic graph in which feature A points to feature B via a directed edge, encoded as a sparse adjacency matrix in the .varp slot of a MuData object^21^, thus naturally supporting N:M mappings. Because the graph is stored as a regular MuData attribute, it requires no changes to the underlying file format and is exposed through a .link namespace extension providing querying, plotting and other utility functions across feature levels.

### Modality-specific quality control and QC metrics

Wherever possible, preprocessing functions are robust to missing values, either by ignoring them (*pp.normalize*) or by modeling them explicitly (*tl.bpca*). Where a method explicitly assumes complete measurements (*pp.scanpy_combat*), AlphaPeptTools detects missing values and raises informative errors. For analyses requiring complete matrices, imputation is matched to the underlying mechanism: MNAR values are modeled with left-shifted Gaussian (*pp.impute_gaussian*), while MAR values can be modeled by k-nearest-neighbor (*pp.impute_knn*), Bayesian principal component analysis^41^ (*pp.impute_bpca*), or simple median imputation (*pp.impute_median*). The metrics module provides measures tailored to proteomics data, including the coefficient of variation (*metrics.coefficient_of_variation*) and pooled median absolute deviation ^42^ (*metrics.pooled_coefficient_of_variation*) to assess quantitative reproducibility during normalization operations, as well as principal component regression^24,25^ (*metrics.principal_component_regression*) to estimate the contribution of covariates to a dataset’s variance. Preprocessing functions additionally document their underlying assumptions and recommended quality-control thresholds in their docstrings, informing users about appropriate cutoffs and how individual steps affect analysis.

### Statistical analysis across feature-levels

AlphaPeptTools’ statistical tools apply uniformly across precursors, peptides, proteins, and genes through the common AnnData interface. For unsupervised analysis, latent embeddings can be computed either by standard principal component analysis (*tl.pca*) on imputed data or by a NaN-aware Bayesian principal-component factor model^41,43^ (*tl.bpca*) and visualized as embeddings (*pl.plot_pca*), factor loadings (*pl.plot_pca_loadings*), and explained variance (*pl.scree_plot*). For differential expression, AlphaPeptTools provides a standard t-test (tl.diff_exp_ttest), a moderated empirical-Bayes t-test for increased power at low replicate numbers^28,44^ (*tl.diff_exp_ebayes*), and proteoform-aware testing via alphaquant^45^ (*tl.diff_exp_alphaquant*). Results from each method are reported in a standardized format and can be visualized through a unified interface (*pl.volcano*).

### Plotting

The ‘pl’ (plotting) module is based on the lightweight, matplotlib-based visualization package *stylia* (https://github.com/ersilia-os/stylia). ‘*pl.create_figure’* generates matplotlib axes with consistent styling, providing an iterator-based way of populating subplots. Complex visualizations can be constructed from basic ‘building-block’-plots through a layering syntax for complex coloring and labelling. We aim to use perceptually uniform and color-vision deficiency (CVD) friendly scientific colors by default^46^. Matplotlib axes objects are always directly accessible, making the syntax especially suitable for LLM-supported analyses, where a basic plot can be extended with custom functionality in a Jupyter notebook.

### Software engineering practices

AlphaPeptTools is developed as an open-source software package, and we ensure software quality via multiple mechanisms: All new code is peer-reviewed by at least one other core maintainer of the package. All modules are automatically and recurrently tested in a continuous integration pipeline, linted (ruff), and type checked (ty), with the core modules (‘iò, ‘pp’, ‘tl’, ‘datà and ‘metrics’) having >80 % unit-test coverage (MuLink: 100% unit-test coverage) to safeguard against regressions. Tutorial notebooks are further automatically tested and serve as integration tests. Documentation is automatically generated and updated via Sphinx based on the scverse cookiecutter template.

### Applications Alzheimer dataset

We analyzed a clinical cohort of 60 Alzheimer’s disease patients (AD)^22^. The respective .raw files were downloaded from the PRIDE repository (PXD016278) and processed with DIA-NN 2.1.0 using a human fasta (UniProt UP000005640, canonical + isoform, release 2022_03; 100,889 entries). The protein dataset was loaded from a DIA-NN report (*io.read_psm_table*), annotated with clinical metadata containing the AD classification (*pp.add_metadata*) and log2-transformed (*pp.nanlog*). Total sample-intensity and number of detected features were assessed via *metrics.calculate_qc_metrics*, followed by visualizing per-sample precursor abundances as well as per-feature median intensity across all protein features, highlighting expected CSF-constituents and hemoglobin contaminants. Then, the dataset was filtered to 75% complete features (*pp.filter_data_completeness*) and missing values imputed from a downshifted normal distribution (*pp.impute_gaussian*). Next, principal components were computed and stored in AnnData’s .obsm attribute (tl.pca)^15^. To assess the tradeoff between batch-correction and retaining biological signal, batch corrected datasets were computed (*pp.scanpy_combat*), and the feature-associated variance quantified using principal component regression (*metrics.principal_component_regression*)^24^ The resulting change in explained variance was visualized as pre-, and post-batch correction barplots (*pl.barplot*). Differential expression analysis between cases and controls was performed via empirical Bayes moderated t-test^28,29^ (*tl.diff_exp_ebayes*), fitting a gene-wise linear model of log-expression on disease condition with sex as an additive covariate.

### Hierarchical feature analysis

In the PELSA chemoproteomics dataset, human cell lysates were treated with either the pan-kinase inhibitor staurosporine or a DMSO control in technical quadruplicates^47^. The respective .raw files were downloaded from the PRIDE repository (PXD034606) and processed with DIA-NN 2.1.0 using a human fasta (UniProt UP000005640, reviewed, canonical + isoform, downloaded 22 July 2025; 42,517 entries). The DIA-NN report was loaded into a precursor AnnData instance (*io.read_psm_table*), annotated with sample metadata (*pp.add_metadata*) and log2-transformed (*pp.nanlog*). Coefficients of variation were calculated on raw counts for each feature in the two treatment levels (*metrics.coefficient_of_variation*). Precursor differential expression was assessed by t-test (*tl.diff_exp_ttest*), empirical Bayes-moderated t-test^28,29^ (*tl.diff_exp_ebayes*) and AlphaQuant^45^ (*tl.diff_exp_alphaquant*), the latter of which operates directly on the raw DIA-NN output. Kinase status was annotated based on www.kinhub.org^48^. To demonstrate working with multiple proteomics data layers, protein and gene-level AnnData instances were created, annotated with metadata, log-transformed and unified with precursor data in a linked MuData instance (*io.mulink_from_anndatas*). The gene MAP3K7 was picked as the top downregulated kinase from the precursor-level differential expression whose gene is associated with at least two protein isoforms. The single regulated precursor of MAP3K7 was highlighted in the gene, protein and precursor volcano plots, as well as in a Boltz-2^49^ predicted structure of MAP3K7 (O43318) with staurosporine, highlighting the binding site proximity of the regulated precursor [Fig2 A].

### scDVP dataset

We analyzed Deep Visual Proteomics (scDVP) data^30^, comprising 598 hepatocytes from 14 healthy donors with 5,175 protein groups measured along the porto-central axis (Download from https://github.com/BorgwardtLab/Human_scDVP_Analysis.git). First, we computed quality control metrics (features detected, detection fraction, total intensity) (*metrics.calculate_qc_metrics*). Next, sample outliers, as defined by more than 2 median absolute deviations (MADs) from the median detected protein count^50^, were removed. Proteins with >30% missing values were dropped (*pp.filter_data_completeness*). Data were log-transformed (*pp.nanlog*) and scaled (*pp.scale_and_center*). Cells with any protein deviating >4 MADs from its median were excluded. PCA (tl.pca) was performed on fully complete proteins, excluding common contaminants (KRT, ALB, DSP, DCD, IGG). To produce Fig. 2B, protein expression was Z-scored per donor, binned into five spatial quantiles, and mean bin-wise profiles computed across donors. TF activity was inferred using decoupler^51^ with the CollecTRI network^52^ and summarized as periportal–pericentral differences. We loaded the available image and shapes data for healthy donor NML10 into the spatialdata framework (v0.7.3,^34^) using the ‘generic’ reader function in spatialdata-io (v0.7.0)^34^, matching measurements on donor ID and sample-wise shape index. Static overlays of tissue images with MS-proteomics measurements were generated with the spatialdata-plot package (v0.4.0)^34^.

### Multimodal non-small cell lung carcinoma (NSCLC) dataset

To perform an integrated multimodal analysis of MS-proteomics, RNA-microarray, and methylation measurements of an NSCLC cohort of 141 patients by Lehtiö et al.^35^, we downloaded the protein group table (gene-symbols_table.txt) from PRIDE (PXD020191), converted it to an AnnData object (*io.read_pg_table*), and preprocessed it as described by the original authors. In short, we filtered for 100% complete features (N=9783; *pp.filter_data_completeness*), log2-transformed raw intensities (*pp.nanlog*), and median centered the data per feature (*pp.scale_and_center*). We loaded preprocessed RNA-data^36^ (118 matched patients) and methylation microarray data^37^ (113 matched patients) from the authors’ supplementary data into AnnData objects. We converted methylation-beta values to M-values^53^ with a pseudocount of 0.01 to approximate a normal distribution. We joined the three AnnData objects into a single MuData object and ran MOFA+^38^ within the muon framework^21^ across the union of samples on scaled data, with 25 factors, Gaussian likelihood, automatic relevance determination and spike-and-slab priors on both factors and weights. We evaluated separation of histological subtypes with Welch’s one-way ANOVA correcting for the multiple testing across 25 factors with the Benjamini Hochberg procedure (FDR<0.01), followed by Games-Howell post-hoc test. To infer modality-wise pathway activities, we ran the decoupler^32^ implementation of gene set enrichment analysis^40^ on the resulting MOFA+ loadings using the hallmarks geneset^54^.

## Extended Data

### Readers: Supported Search Engines for IO

**Table 1.** Supported search engine outputs for scverse-compatible analysis packages.

|  | DIA/DDA | Table Format | AlphaPeptTools<br>(v0.3.0) | msmu<sup>1</sup><sub>8</sub><br>(v0.3.1) | Proteopy<sup>1</sup><sub>7</sub> (v0.1.1) | protdat<sub>a</sub><br>(v0.1.4) | OpenDVP<sup>1</sup><sub>9</sub><br>(v0.7.3) |
| --- | --- | --- | --- | --- | --- | --- | --- |
| **alphadia** | DIA | PSM | Yes | - | - | - | - |
|  |  | PG | Yes | - | - | - | - |
| **alphapept** | DDA | PSM | Yes | - | - | - | - |
|  |  | PG | Yes | - | - | - | - |
| **DIA-NN** | DIA | PG | Yes | - | - | Yes | Yes |
|  |  | PSM | Yes | Yes | Yes | - | - |
| **MSFragger +**<br>**FragPipe** | DIA/DDA | PG | Yes | - | - | Yes | - |
|  |  | PSM | Yes | Yes<br>(only DDA) | - | - | - |
| **Spectronaut** | DIA | PG | Yes | - | - | Yes | - |
|  |  | PSM | Yes | - | - | - | - |
| **MaxQuant** | DDA +<br>DIA | PG | Yes | - | - | Yes | - |
|  |  | PSM | Yes | Yes<br>(only DDA) | - | - | - |
| **Perseus** | agnostic | (processed ) | No | - | - | - | Yes |
| **Pfind** | DDA | PG | No | - | - | - | - |
|  |  | PSM | Yes | - | - | - | - |
| **Sage** | DDA | PG | No | - | - | - | - |
|  |  | PSM | Yes | Yes | - | - | - |
| **Delpi** | DIA | PSM | - | Yes | - | - | - |
| **Proteome**<br>**Discoverer** | DDA | PG | Yes | - | - | - | - |
| **mztab** | agnostic | PG | Yes | - | - | Yes | - |
| **directlfq** | agnostic | (PG,<br>processed) | Yes | - | - | - | - |
| **Complexity:**<br>**User-facing**<br>**reader**<br>**functions** |  |  | 2 | 5 | 2 (+1<br>generic) | 4 | 2 |
| **Standardize**<br>**d Column**<br>**Output** |  |  | Yes | No | No |  | No |

### Features

**Table 2.** Supported features for scverse-compatible analysis packages.

| Feature |  | AlphaPeptTools<br>(v0.3.0) | msmu<sup>18</sup><br>(v0.3.1) | scanpy<sup>15</sup><br>(v1.12.3) | proteopy<sup>17</sup><br>(v0.1.1) | OpenDVP<sup>19</sup><br>(v0.7.3) |
| --- | --- | --- | --- | --- | --- | --- |
| **Robustness**<br>**against NaNs** |  | Yes | Yes | No | Yes | Yes |
| **Hierarchical**<br>**Feature**<br>**Representation** |  | Yes. Multi-<br>layered<br>(MuData) +<br>Explicit feature<br>connections<br>(MuLink) | Yes. Multi-<br>layered<br>(MuData) | No | Yes. Custom<br>indexing | No |
| **Imputation** | **MAR** | Yes | No | No | No | No |
|  | **MNAR** | Yes | No | No | Yes | Yes |
| **Batch correction** |  | Yes (Combat,<br>IRS) | Yes (Combat,<br>IRS, Median<br>Centering,<br>LOWESS) | Yes<br>(Combat) | No | No |
| **Dimensionality**<br>**Reduction** | **General** | Yes (PCA) | Yes (PCA;<br>UMAP) | Yes (PCA;<br>UMAP;<br>tSNE) | No | No |
|  | **Nan-**<br>**aware** | Yes (BPCA) | No | No | No | No |
| **Differential**<br>**expression** |  | Yes (t-test,<br>welch,<br>moderated t-test) | Yes (t-test,<br>welch,<br>Wilcoxon,<br>Moderated t-test,<br>permutation) | Yes (t-test,<br>wilcoxon) | Yes (t-test,<br>welch,<br>anova) | Yes (t-test,<br>anova,<br>bootstrap) |
| **Isoform aware**<br>**differential**<br>**expression** |  | Yes (alphaquant) | No | No | Yes (COPF) | No |
| **Feature**<br>**aggregation** |  | No (search<br>engine based) | Yes | No | Yes | No |

## Appendix

### Code Availability

AlphaPeptTools is available under an open Apache 2.0 license on GitHub:

- https://github.com/MannLabs/alphapepttools.git

Documentation, including instructive notebooks to reproduce the analysis of this manuscript is available at

- https://mannlabs.github.io/alphapepttools

The MuLink package is available under an open MIT license on GitHub:

- https://github.com/lucas-diedrich/mulink.git

with documentation being available under

- https://mulink.readthedocs.io

Code and raw data to reproduce the figures in this manuscript is deposited on GitHub and Figshare:

- https://github.com/MannLabs/alphapepttools-manuscript.git
- https://doi.org/10.6084/m9.figshare.33514423

## Data Availability

All data to reproduce the performed analyses is available via the alphapepttools.data.get_data function and deposited on Figshare https://doi.org/10.6084/m9.figshare.33514423.

## Acknowledgements

We thank Ilia Kats for his helpful comments on the integration of MuLink with MuData. This study was supported by the Max Planck Society for Advancement of Science.

## Use of generative AI

Generative AI tools (Claude Opus 4.8, 5.0, 5.1) were used to assist software development and manuscript preparation. All AI-generated suggestions were carefully reviewed, edited, and validated by the authors. The authors take full responsibility for the content of the software and manuscript.

## Notes

https://github.com/MannLabs/AlphaPeptTools.git

### Competing Interest Statement

The authors have declared no competing interest.

